# *%svy_freqs:* A generic SAS macro for cross-tabulation between a factor and a by-group variable given a third variable and creating publication-quality tables using data from complex surveys

**DOI:** 10.1101/771303

**Authors:** Jacques Muthusi, Samuel Mwalili, Peter Young

**Author notes:** **Corresponding author:** Email address (J. Muthusi).

## Abstract

**Introduction:** In epidemiological studies, cross-tabulations are a simple but important tool for understanding the distribution of socio-demographic characteristics among study participants. They become more useful when comparisons are presented using a by-group variable such as key demographic characteristic or an outcome status; for instance, sex or the presence or absence of a disease status. Most available statistical analysis software can easily perform cross-tabulations, however, output from these must be processed further to make it readily available for review and use in a publication. In addition, performing three-way cross-tabulations of complex survey data such as those required to show the distribution of disease prevalence across multiple factors and a by-group variable is not easily implemented directly using available standard procedures of commonly used statistical software.

**Methods:** We developed a generic SAS macro, ***%svy_freqs,*** to create quality publication-ready tables from cross-tabulations between a factor and a by-group variable given a third variable using survey or non-survey data. The SAS macro also performs classical two-way cross-tabulations and refines output into publication-quality tables. It provides extra features not available in existing procedures such as ability to incorporate parameters for survey design and replication-based variance estimation methods, performing validation checks for input parameters, transparently formatting character variable values into numeric ones and allowing for generalizability.

**Results:** We demonstrate the application of the SAS macro in the analysis of data from the 2013-2014 National Health and Nutrition Examination Survey (NHANES), a complex survey designed to assess the health and nutritional status of adults and children in the United States (U.S.).

**Conclusion:** The SAS code use to develop the macro is simple yet comprehensive, easy to follow, straightforward for the end user and simple for a SAS programmer to extend. The SAS macro has shown to shorten turn-around time for statistical analysis, eliminate errors when preparing output, and support reproducible research.

## Introduction

Cross-tabulations are a basic but important tool for understanding the distribution of socio-demographic characteristics among study or survey participants in the fields of epidemiology and disease surveillance. They are handy especially where multiple tables are required. This is very useful especially when comparisons need to be performed separately by a by-group variable such as a key demographic characteristic, e.g. sex, or an outcome status such as positive or negative test result for a disease. Cross-tabulations can be even more informative if one is interested in distribution of disease prevalence among selected factor variables (table rows) and a by-group variable (table columns). This is useful in cases where the association between disease prevalence and risk factors or exposures needs to be stratified, for instance, by sex or geographic region.

Almost all available statistical analysis software can easily perform cross-tabulations, however, output from these must be processed further to make them readily available for review and use in a publication. In Stata, one can use `table, tabulate` [1] commands or Stata user’s community-contributed programs like `tabout` [2] or `tabmult` [3]. In SAS, there exist a limited number of commands or macros for creating publication-quality tables [4–9] but they suffer from limitations of flexibility, usability and generalizability. For instance, Sunesara et al. [4], provide a SAS macro that works with only a two-level/binary by-group variable which must further be manually re-coded as 0 or 1, hence making it less flexible and more difficult to use with three or more levels of a by-group variable. Though the Sunesara macro allows users to specify survey design variables, it does not provide for domain analysis and output must be processed further to be useful. For instance, variable labels, which are more meaningful, are used instead of variable names and usually appear in the same column above the corresponding categories to match layout for epidemiological publications. The macro further provides only means for continuous variables, rather than an option for robust measure of central tendency medians.

Other SAS macros, including *%YAMGAST*, by Xiong [5] and *%SummaryTable* by Zhouet al. [7] have been proposed for use in the pharmaceutical industry and not in epidemiology settings where survey design should be incorporated during analysis. Zuo [8] proposed a set of disparate SAS macros for performing cross-tabulation. However, for the analyst to be able to use these correctly they require knowing the order in which they flow and they require specifying the same input parameters repeatedly. This is can be inefficient and can introduce several uncertainties leading to errors during the analysis. Martin [6] further provided some useful guidelines for making descriptive summary tables that are ready to review. In addition, the SAS macros available do not provide the analyst with options for specifying replication-based variance estimation methods including Jackknife (JK) or Balanced Repeated Replication (BRR). These methods are crucial in obtaining correct variances for survey estimates in presence of survey non-response, hence providing valid variance estimates.

We have developed a SAS macro which overcomes the described shortcomings while promoting reproducible research principles [8] such as transparency, reproducibility and reusability, which are attracting detailed attention in epidemiological research [10–14]. It further provides for replication-based variance estimation methods as well as enforce validation checks for input parameters.

## Methods

### Sample survey methods

Sample surveys are often used to study a population using a sample instead of studying the entire population. Simple or complex survey methods can be used to select the sample. [15–20] discuss in details the theory and application of various sample survey methods including simple random sampling, stratified sampling, clustered sampling, and multi-stage sampling among others.

### The *%svy_freqs* SAS macro

In SAS software, cross-tabulations can be performed using the FREQ or SURVEYFREQ procedures for categorical variables and the MEAN or SURVEYMEANS procedures for continuous variables [21]. The SAS macro, ****%svy_freqs****, written in SAS software version 9.3 [22], uses the SURVEYFREQ and SURVEYMEANS procedures to perform the cross-tabulation and output frequencies, totals and percentages. The macro runs sequentially starting with cross-tabulations for categorical variables followed by continuous variables. All output is then processed using the TEMPLATE, PROC REPORT procedures and the output delivery system (ODS) to create a publication-quality table, similar to a typical Table 1 or Table 2 of a manuscript.

**Table 1:**
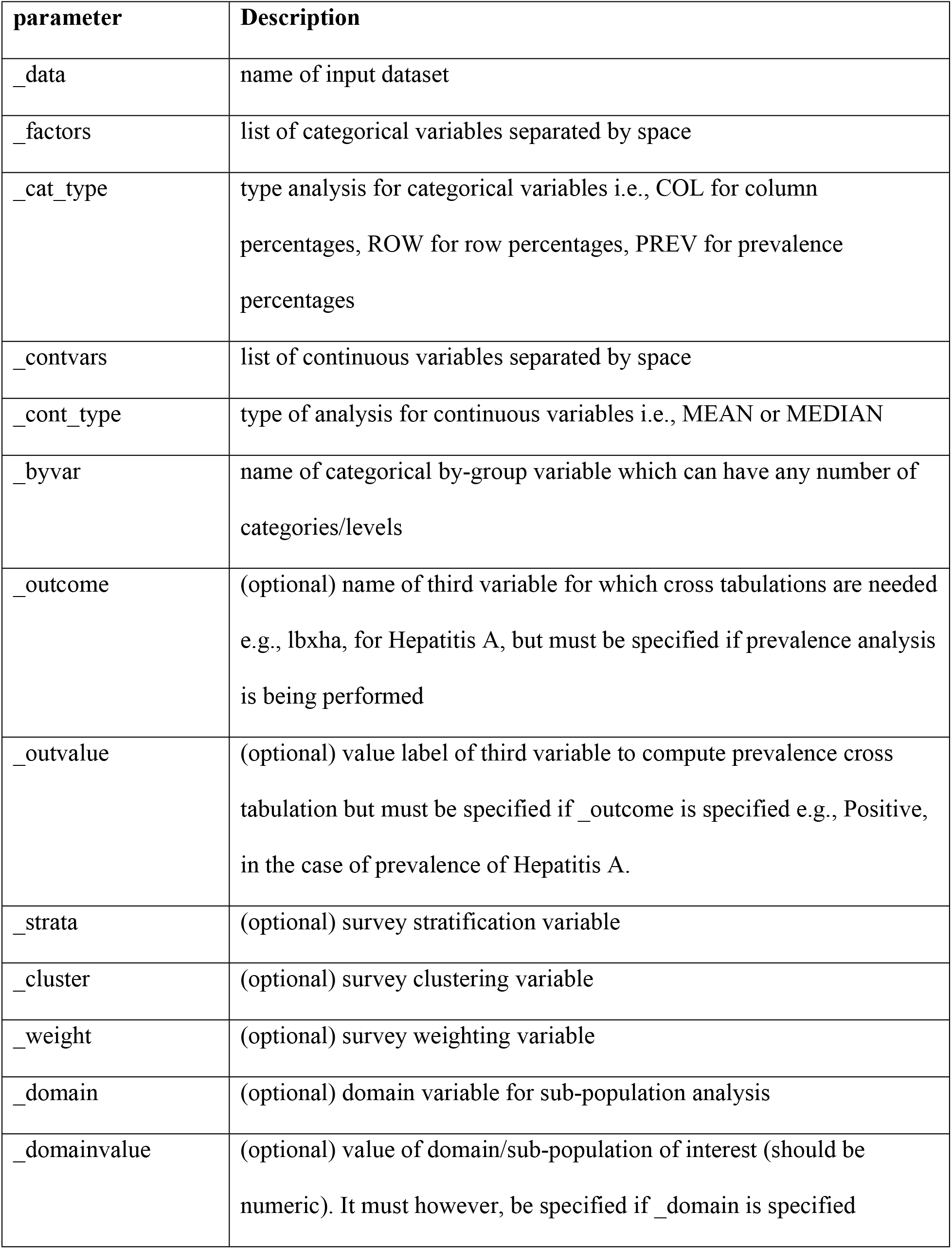
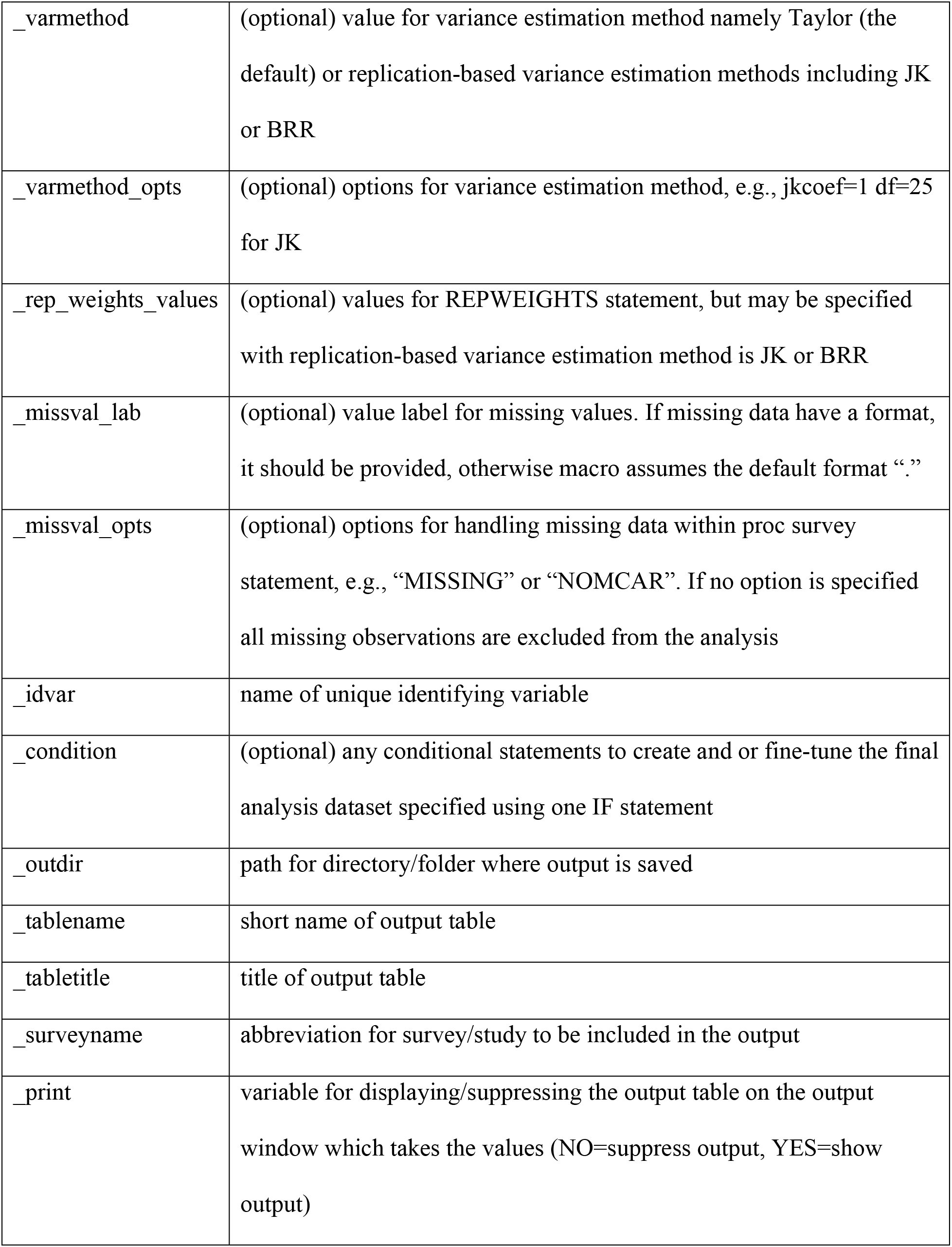
Input parameters for *%syy_freqs* **macro**.

| **parameter** | **Description** |
| --- | --- |
| _data | name of input dataset |
| _factors | list of categorical variables separated by space |
| _cat_type | type analysis for categorical variables i.e., COL for column percentages, ROW for row percentages, PREV for prevalence percentages |
| _contvars | list of continuous variables separated by space |
| _cont_type | type of analysis for continuous variables i.e., MEAN or MEDIAN |
| _byvar | name of categorical by-group variable which can have any number of categories/levels |
| _outcome | (optional) name of third variable for which cross tabulations are needed e.g., lbxha, for Hepatitis A, but must be specified if prevalence analysis is being performed |
| _outvalue | (optional) value label of third variable to compute prevalence cross tabulation but must be specified if _outcome is specified e.g., Positive, in the case of prevalence of Hepatitis A. |
| _strata | (optional) survey stratification variable |
| _cluster | (optional) survey clustering variable |
| _weight | (optional) survey weighting variable |
| _domain | (optional) domain variable for sub-population analysis |
| _domainvalue | (optional) value of domain/sub-population of interest (should be numeric). It must however, be specified if _domain is specified |
| _varmethod | (optional) value for variance estimation method namely Taylor (the default) or replication-based variance estimation methods including JK or BRR |
| _varmethod_opts | (optional) options for variance estimation method, e.g., jkcoef=1 df=25 for JK |
| _rep_weights_values | (optional) values for REPWEIGHTS statement, but may be specified with replication-based variance estimation method is JK or BRR |
| _missval_lab | (optional) value label for missing values. If missing data have a format, it should be provided, otherwise macro assumes the default format “.” |
| _missval_opts | (optional) options for handling missing data within proc survey statement, e.g., “MISSING” or “NOMCAR”. If no option is specified all missing observations are excluded from the analysis |
| _idvar | name of unique identifying variable |
| _condition | (optional) any conditional statements to create and or fine-tune the final analysis dataset specified using one IF statement |
| _outdir | path for directory/folder where output is saved |
| _tablename | short name of output table |
| _tabletitle | title of output table |
| _surveyname | abbreviation for survey/study to be included in the output |
| _print | variable for displaying/suppressing the output table on the output window which takes the values (NO=suppress output, YES=show output) |

**Table 2:** Participants socio-demographic characteristics by sex (Col %), N=522 ^π^.

| Characteristic <sup>£</sup> | Male |  |  | Female |  |  | Total <sup>€</sup> |  |  |
| --- | --- | --- | --- | --- | --- | --- | --- | --- | --- |
| | Unweighted n <sup>@</sup> | Weighted % (or median) <sup>¥</sup> | 95 % CI (or IQR) <sup>\$</sup> | Unweighted n | Weighted % (or median) <sup>¥</sup> | 95 % CI (or IQR) <sup>\$</sup> | Unweighted N | Weighted % (or median) <sup>¥</sup> | 95 % CI (or IQR) <sup>\$</sup> |
| Age in years at screening |  |  |  |  |  |  |  |  |  |
| 20-39 | 46 | 12.7 | (7.9 - 17.5) | 6 | 19.2 | (2.8 - 35.6) | 52 | 13.3 | (8.6 - 18.0) |
| 40-59 | 114 | 25.2 | (19.9 - 30.5) | 24 | 61.8 | (38.2 - 85.3) | 138 | 28.3 | (21.4 - 35.2) |
| >= 60 | 326 | 62.1 | (56.0 - 68.1) | 6 | 19.1 | (2.2 - 35.9) | 332 | 58.4 | (51.5 - 65.4) |
| Total | 486 | 100 | ( - ) | 36 | 100 | ( - ) | 522 | 100 | ( - ) |
| Race/Hispanic origin |  |  |  |  |  |  |  |  |  |
| Mexican American | 23 | 2.6 | (0.7 - 4.5) | 0 | . | (. - .) | 23 | 2.4 | (0.7 - 4.1) |

|  | Male |  |  | Female |  |  | Total € |  |  |
| --- | --- | --- | --- | --- | --- | --- | --- | --- | --- |
| Characteristic £ | Unweighted n @ | Weighted % (or median) ¥ | 95 % CI (or IQ R) \$ | Unweighted n | Weighted % (or median) | 95 % CI (or IQ R) | Unweighted N | Weighted % (or median) | 95 % CI (or IQ R) |
| Other Hispanic | 25 | 3.2 | (1.6 - 4.8) | 2 | 2.9 | (0.0 - 7.5) | 27 | 3.2 | (1.5 - 4.8) |
| Non-Hispanic White | 296 | 80.1 | (74.0 - 86.2) | 16 | 68.3 | (52.4 - 84.3) | 312 | 79.1 | (73.1 - 85.2) |
| Non-Hispanic Black | 115 | 10.2 | (6.4 - 14.0) | 17 | 27.7 | (14.3 - 41.1) | 132 | 11.6 | (7.8 - 15.5) |
| Other Race | 27 | 3.9 | (1.5 - 6.2) | 1 | 1.1 | (0.0 - 3.6) | 28 | 3.6 | (1.5 - 5.8) |
| Total | 486 | 100 | ( - ) | 36 | 100 | ( - ) | 522 | 100 | ( - ) |
| Served in a foreign country |  |  |  |  |  |  |  |  |  |
| Yes | 261 | 52.4 | (46.2 - 58.6) | 14 | 36.8 | (18.7 - 54.9) | 275 | 51.1 | (45.7 - 56.5) |
| No | 224 | 47.6 | (41.4 - 53.8) | 22 | 63.2 | (45.1 - 81.3) | 246 | 48.9 | (43.5 - 54.3) |
| Missing | 1 | 0.6 | (0.0 - 2.0) | 0 | . | (. - .) | 1 | 0.6 | (0.0 - 1.8) |
| Total | 486 | 100 | ( - ) | 36 | 100 | ( - ) | 522 | 100 | ( - ) |
| Hepatitis A antibody |  |  |  |  |  |  |  |  |  |
| Positive | 205 | 37.7 | (34.0 - 41.4) | 15 | 39.7 | (18.0 - 61.3) | 220 | 37.9 | (34.8 - 40.9) |
| Negative | 268 | 62.3 | (58.6 - 66.0) | 20 | 60.3 | (38.7 - 82.0) | 288 | 62.1 | (59.1 - 65.2) |
| Characteristic £ | Unweighted n @ | Weighted % (or median) ¥ | 95 % CI (or IQR) \$ | Unweighted n | Weighted % (or median) | 95 % CI (or IQR) | Unweighted N | Weighted % (or median) | 95 % CI (or IQR) |
| Missing | 13 | 1.7 | (0.7 - 2.7) | 1 | 2.1 | (0.0 - 6.0) | 14 | 1.7 | (0.8 - 2.7) |
| Total | 486 | 100 | ( - ) | 36 | 100 | ( - ) | 522 | 100 | ( - ) |
| Age in years at screening | 486 | 64.2 | (51.3 - 73.0) | 36 | 50.3 | (41.1 - 53.8) | 522 | 63.5 | (49.5 - 72.2) |

π=Analysis domain sample size
€=By-group variable
£= column for listing factor variables labels and corresponding categories
@=column for listing raw frequencies
¥=column for listing weighted column/row/prevalence % (for categorical factors) or median/mean (for continuous factors)
\$=column for weighted 95% confidence interval (for column/row/prevalence/mean % estimates) or interquartile range, IQR (for median estimates)

The macro is composed of seven sub-macros, which are called within the main macro. The first three sub-macros, ***%svy_prev***, ***%svy_row***, ***%svy_col,*** perform cross tabulation for categorical variables. The sub-macros ***%svy_mean*** and ***%svy_median*** provide cross tabulations for continuous variables by computing the mean or median, respectively, whereas ***%charvar, %distcolval*** and ***%runquit*** are used for internal processing. The sub-macro, ***%svy_prev***, which provides prevalence percentages, performs three-way cross-tabulations between a factor and a by-group variable given a third variable. The `_outcome` and `_outvalue`, which are the parameters for which prevalence is to be computed, must be specified. Analysis type, `_cat_type`, must be specified as equal to `PREV`. If not specified, the macro automatically generates a new variable, `_freq` whose value equals 1 for all study subjects in the analysis dataset, and proceeds with the analysis as though it were for two-way cross-tabulations with row percentages, similar to the ***%svy_row*** sub-macro. The `_outcome` and `_outvalue` parameters may be omitted for two-way cross-tabulations performed using ***%svy_row*** (for row percentages), ***%svy_col*** (for column percentages) sub-macros. The ***%charvar*** macro encodes variables with character values to numeric ones for successful execution by the ***%svy_freqs*** macro. The ***%distcolval*** macro is used to format output by putting the variable label of a factor in the same column as the factor levels hence providing a natural display of results as often expected in epidemiological publications and reports. The ***%runquit*** macro enforces in-built SAS validation checks on input parameters and tests for logical errors. It halts the macro from execution and prints out the error on the log window for the user to address. The user should specify input parameters to the main macro that are described in Table 1 unless the description is prefixed by (optional). The user, however, does not interact with the sub-macros. To achieve full potential of the SAS macro, the user must ensure that the analysis dataset is clean, analysis variables are well labelled, and values of variables have been converted into appropriate SAS formats before they can be input to the macro call.

The SAS macro, ***%svy_freqs***, was specially developed for conducting three-way cross-tabulation such as those required in showing the distribution of disease prevalence across multiple factor variables and a by-group variable. However, two-way cross tabulations between factors and by-group variables that are commonly used in epidemiological studies or surveys are also performed by the macro. For instance, if users are interested in showing distribution of study participants by a given by group variable, then column percentages which are most appropriate are obtained using the `COL` option. If the by-group variable is an outcome of interest such as positive or negative diagnostic test results, then the `row` percentages are most appropriate and can obtained using the ROW option. The by-group variable can have more than two categories which can be encoded as either a numeric or character variable. For the distribution of continuous variables, one can specify the type of statistics to compute i.e., either mean or median.

Where the data to be analyzed come from a complex survey, our macro allows users to specify study design variables containing strata, cluster, and design weights. The macro also provides for domain analysis for sub-populations which can be specified using the `_domain` and `_domainvalue` parameters. Variance estimation methods for complex surveys including the Taylor series or the replication based methods such as JK and BRR can be specified in the macro using the `_varmethod` parameter. Where JK or BRR methods are specified, users may also specify values for REPWEIGHTS statement using the `_rep_weights_values` parameter. To customized the replication process, users should add options via the `varmethod_opts` parameter. [16, 21, 23] provides detailed description of the use of replication methods for variance estimating in sample surveys. If variance estimation method is not specified then the macro uses the default Taylor Series.

Complex surveys are usually met with survey non-response. The macro has also been developed to handle missing data accordingly. `_missval_lab` parameter allows user to explicitly identify the value label for missing data. If not specified the macro assumes the default value “.”. The `_missval_opts` parameter specifies options of how missing data are handled. Acceptable options are “MISSING” and “NOMCAR”. If this option is not specified then all missing values are excluded from the analysis. [21, 24–26] provides detailed description on how to use various missing data options. Data from non-survey settings are analyzed by leaving the survey-design parameters unspecified.

If the analysis includes non-coded character variables, the macro automatically encodes them into numeric ones prior to analysis. The macro further provides natural display of results from epidemiological surveys by processing the final output into a refined publication-quality table, which is output into word processing and spreadsheet programs for immediate use in publications or for additional formatting if needed. The macro becomes handy when numerous tables with similar structure need to be generated such as those shown in the Kenya AIDS Indicator Surveys (KAIS) [27, 28] and the Kenya Demographic Health Surveys [29–31]. Its use in the analysis of data from these types of surveys could significantly shorten the analysis period while supporting generation of high-quality outputs.

## Results

### Example of macro call to analyze the NHANES dataset

We demonstrate the application of SAS macro, ***%svy_freqs,*** in the analysis of a dataset from the 2013-2014 National Health and Nutrition Examination Survey (NHANES). NHANES is complex surveys designed to assess the health and nutritional status of adults and children in the United States (U.S.). A detailed description of the survey design and contents is available elsewhere [32]. The NHANES dataset [33] is publicly available online for free from the U.S. Centers for Disease Control and Prevention (CDC) at: https://www.cdc.gov/nchs/nhanes/Index.htm.

The dataset (`clean_nhanes`) used for this analysis comprised of the following variables: `riagendr` (gender), `ridageyr` (age in years at screening), `ridreth1,` (race/Hispanic origin), `dmqadfc` (service in a foreign country), `dmdeduc2` (education level among adults aged 20+ years), and `dmdmartl` (marital status). We used the macro to generate three different tables with the main one (Table 4 with prevalence percentages) showing the distribution of hepatitis A prevalence across selected socio-demographic characteristics and by sex. The next tables show the distribution of participants’ socio-demographic characteristics by sex (Table 2 with column percentages) and by hepatitis A antibody test result (Table 3 with row percentages). The primary outcome considered was the binary variable `lbxha` (Hepatitis A antibody test result). The aim of the analysis was to show the distribution of hepatitis A among participants aged 20+ years who had served active duty in the U.S. Armed Forces (`dmqmiliz`). We also show participant’s socio-demographic characteristics by sex and by hepatitis A antibody test result. Appropriate survey weights `wtmec2yr` (sample weights for participants with a medical examination) were applied. The working denominator was N=542. However, 180 observations were dropped during analysis because they had non-positive weights. In addition, analysis domain sample size was calculated and added at the end of each table title.

**Table 3:**
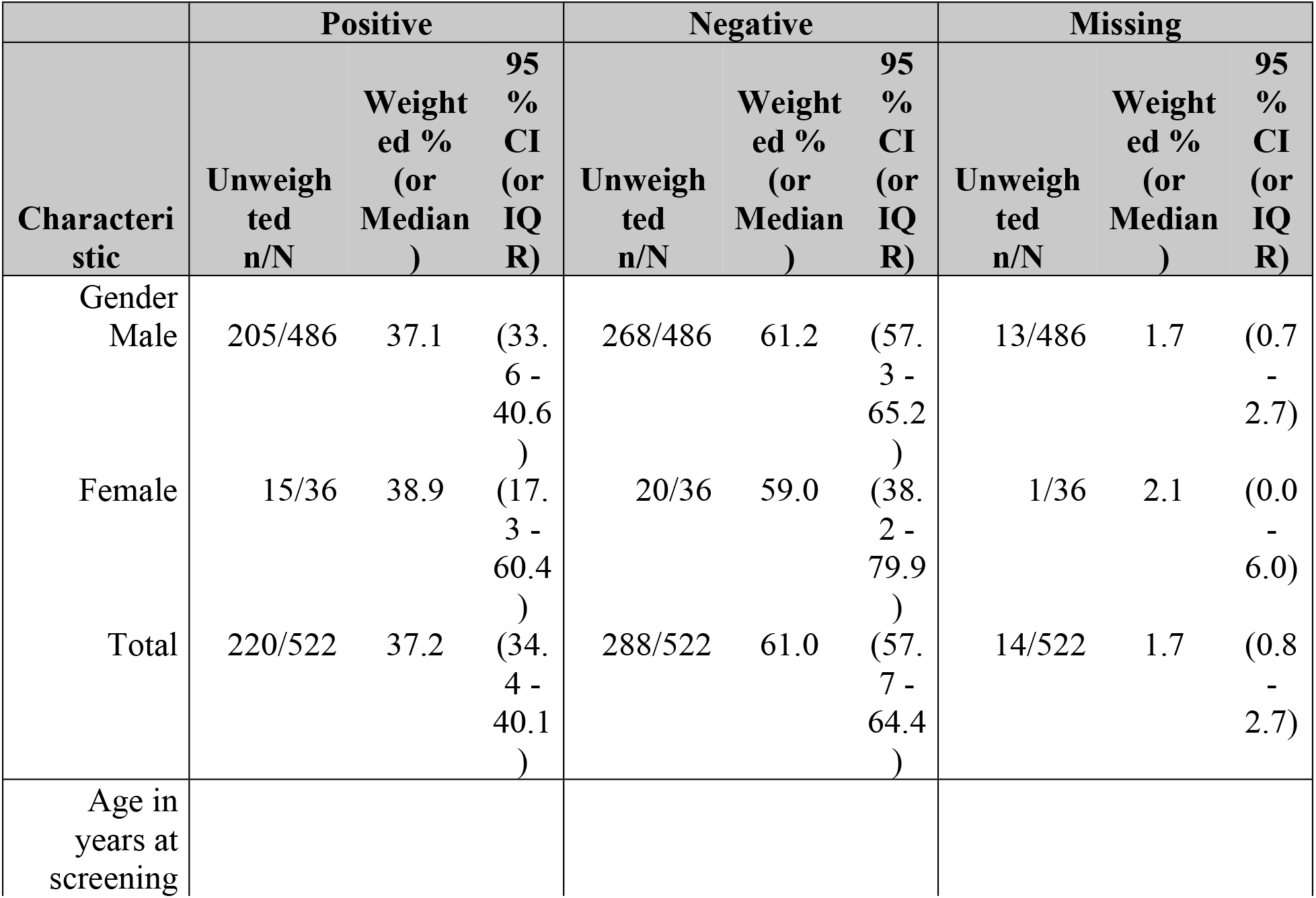
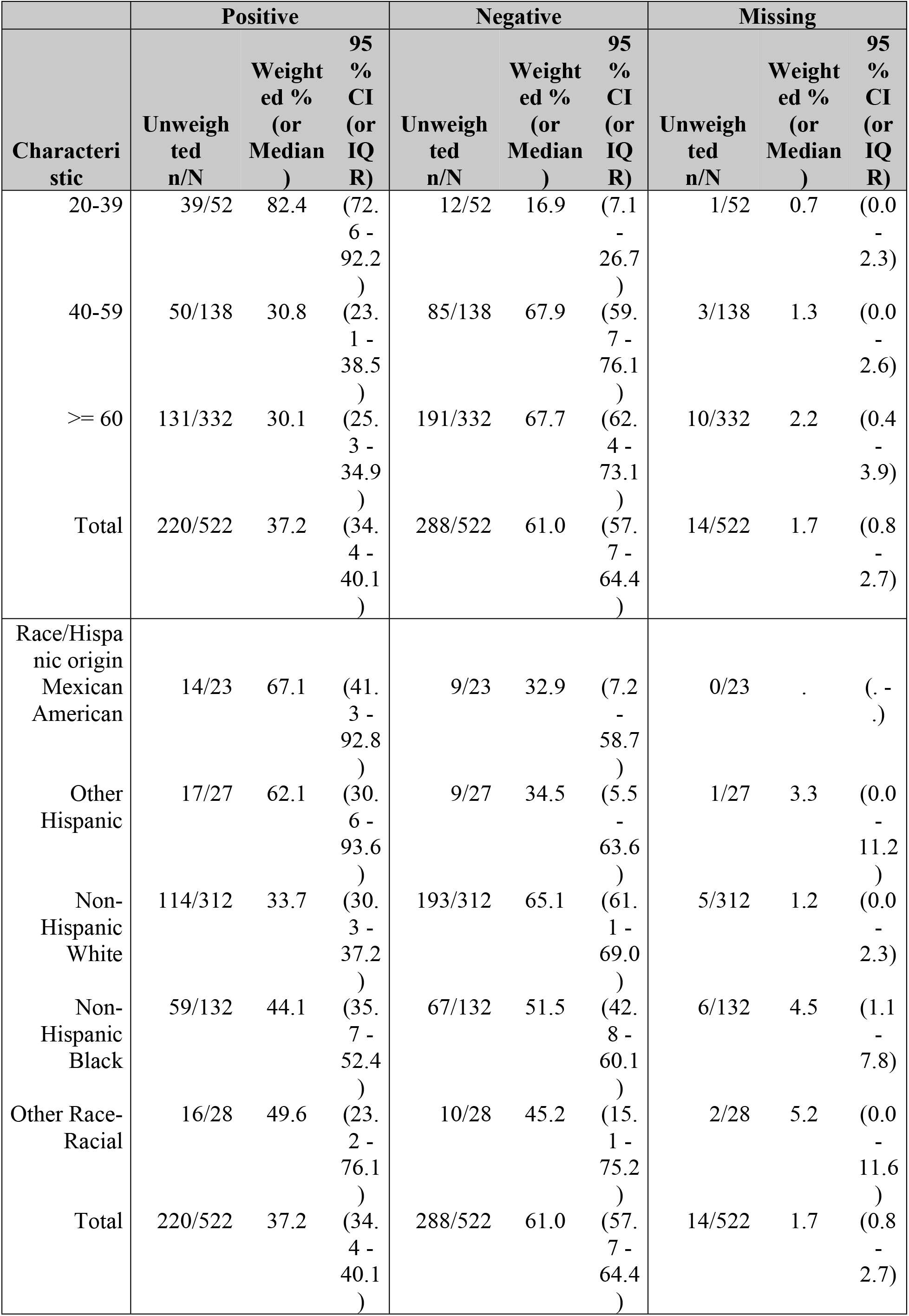
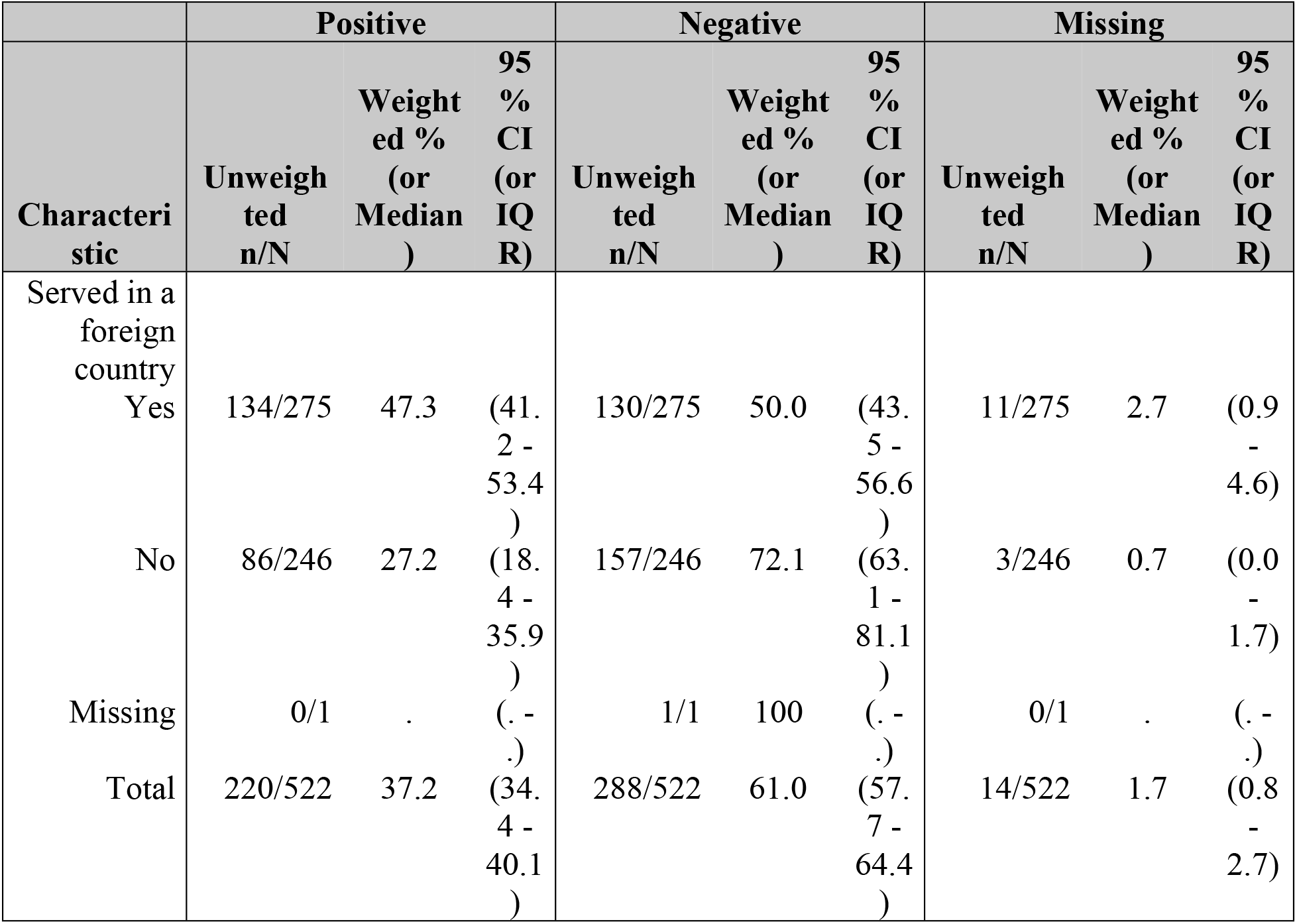
Socio-demographic characteristics by Hepatitis A status (Row %), N=522.

**Table 4:**
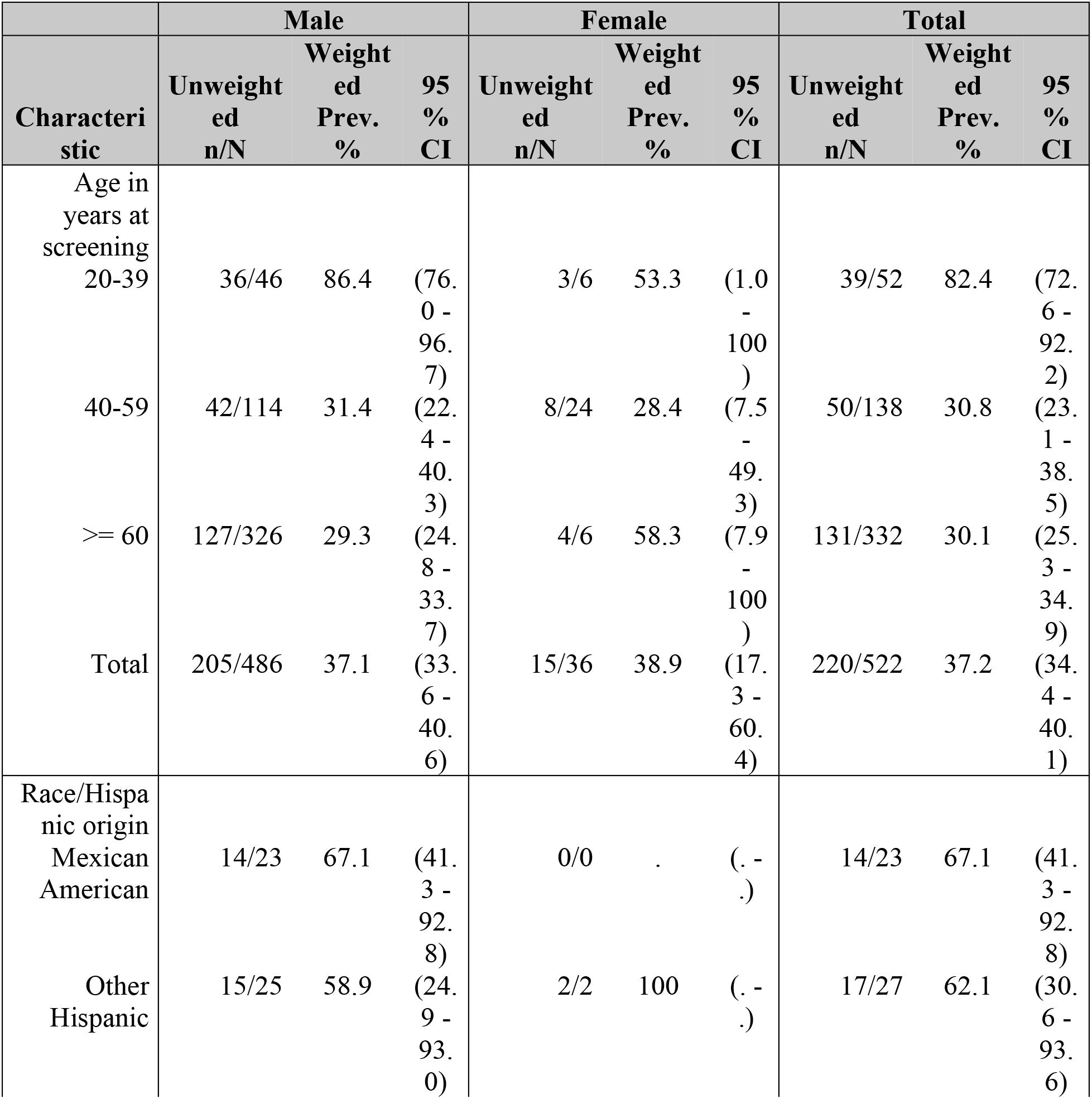
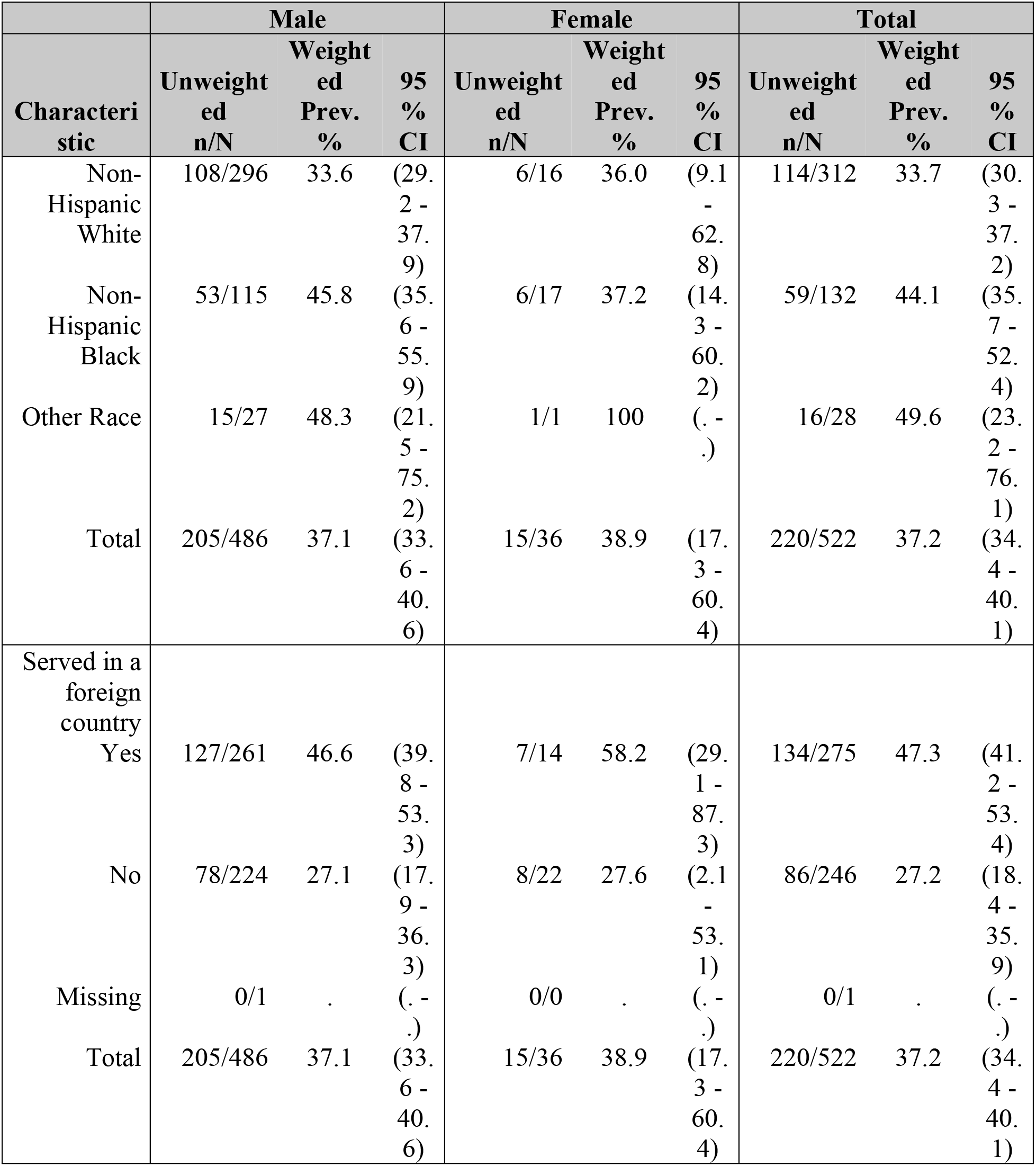
Distribution of Hepatitis A prevalence by selected socio-demographic characteristics and sex (Prevalence %), N=522.

|  | Male |  |  | Female |  |  | Total |  |  |
| --- | --- | --- | --- | --- | --- | --- | --- | --- | --- |
| Characteri<br>stic | Unweight<br>ed<br>n/N | Weight<br>ed<br>Prev.<br>% | 95<br>%<br>CI | Unweight<br>ed<br>n/N | Weight<br>ed<br>Prev.<br>% | 95<br>%<br>CI | Unweight<br>ed<br>n/N | Weight<br>ed<br>Prev.<br>% | 95<br>%<br>CI |
| Age in<br>years at<br>screening |  |  |  |  |  |  |  |  |  |
| 20-39 | 36/46 | 86.4 | (76.<br>0 -<br>96.<br>7) | 3/6 | 53.3 | (1.0<br>-<br>100<br>) | 39/52 | 82.4 | (72.<br>6 -<br>92.<br>2) |
| 40-59 | 42/114 | 31.4 | (22.<br>4 -<br>40.<br>3) | 8/24 | 28.4 | (7.5<br>-<br>49.<br>3) | 50/138 | 30.8 | (23.<br>1 -<br>38.<br>5) |
| >= 60 | 127/326 | 29.3 | (24.<br>8 -<br>33.<br>7) | 4/6 | 58.3 | (7.9<br>-<br>100<br>) | 131/332 | 30.1 | (25.<br>3 -<br>34.<br>9) |
| Total | 205/486 | 37.1 | (33.<br>6 -<br>40.<br>6) | 15/36 | 38.9 | (17.<br>3 -<br>60.<br>4) | 220/522 | 37.2 | (34.<br>4 -<br>40.<br>1) |
| Race/Hispa<br>nic origin |  |  |  |  |  |  |  |  |  |
| Mexican<br>American | 14/23 | 67.1 | (41.<br>3 -<br>92.<br>8) | 0/0 | . | (. -<br>) | 14/23 | 67.1 | (41.<br>3 -<br>92.<br>8) |
| Other<br>Hispanic | 15/25 | 58.9 | (24.<br>9 -<br>93.<br>0) | 2/2 | 100 | (. -<br>) | 17/27 | 62.1 | (30.<br>6 -<br>93.<br>6) |
| Characteristic | Unweighted<br>n/N | Weighted<br>Prev.<br>% | 95<br>%<br>CI | Unweighted<br>n/N | Weighted<br>Prev.<br>% | 95<br>%<br>CI | Unweighted<br>n/N | Weighted<br>Prev.<br>% | 95<br>%<br>CI |
| Non-Hispanic White | 108/296 | 33.6 | (29.2 - 37.9) | 6/16 | 36.0 | (9.1 - 62.8) | 114/312 | 33.7 | (30.3 - 37.2) |
| Non-Hispanic Black | 53/115 | 45.8 | (35.6 - 55.9) | 6/17 | 37.2 | (14.3 - 60.2) | 59/132 | 44.1 | (35.7 - 52.4) |
| Other Race | 15/27 | 48.3 | (21.5 - 75.2) | 1/1 | 100 | (. - .) | 16/28 | 49.6 | (23.2 - 76.1) |
| Total | 205/486 | 37.1 | (33.6 - 40.6) | 15/36 | 38.9 | (17.3 - 60.4) | 220/522 | 37.2 | (34.4 - 40.1) |
| Served in a foreign country |  |  |  |  |  |  |  |  |  |
| Yes | 127/261 | 46.6 | (39.8 - 53.3) | 7/14 | 58.2 | (29.1 - 87.3) | 134/275 | 47.3 | (41.2 - 53.4) |
| No | 78/224 | 27.1 | (17.9 - 36.3) | 8/22 | 27.6 | (2.1 - 53.1) | 86/246 | 27.2 | (18.4 - 35.9) |
| Missing | 0/1 | . | (. - .) | 0/0 | . | (. - .) | 0/1 | . | (. - .) |
| Total | 205/486 | 37.1 | (33.6 - 40.6) | 15/36 | 38.9 | (17.3 - 60.4) | 220/522 | 37.2 | (34.4 - 40.1) |

The macros were run sequentially after specifying required parameters as shown in Fig 1-3. The results presented here are for purely for illustrative purposes only and do not follow from any specific survey objective. Readers should consult the NHANES analytic guidelines on variable definitions, analytical and statistical recommendations that are available online at https://wwwn.cdc.gov/nchs/nhanes/analyticguidelines.aspx

**Fig 1.**
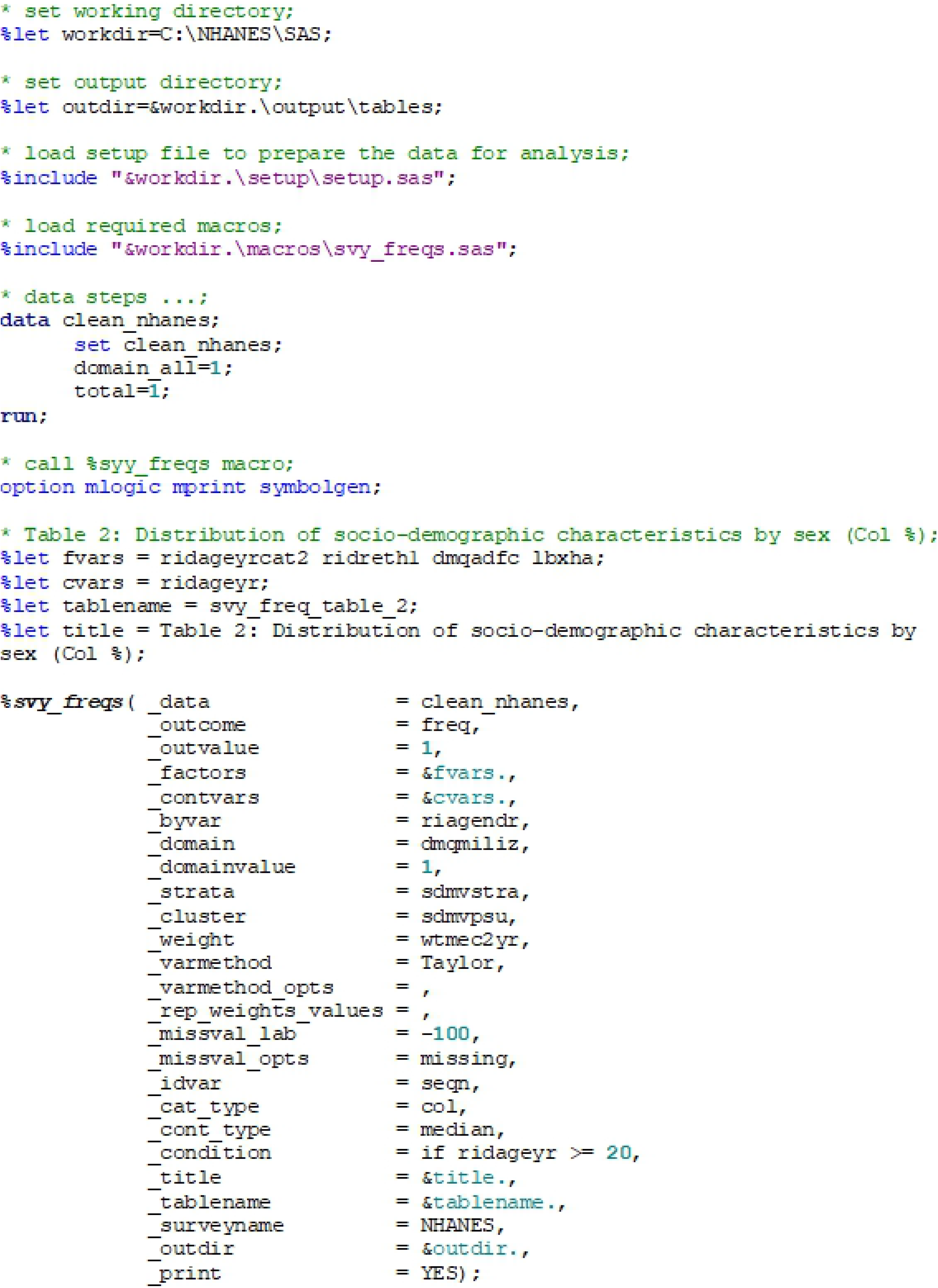
Sample *%svy_freqs* macro call to output column percentages.

The SAS output from the macro consists of several tables specifically for holding parameter estimates, corresponding 95% CI for percentages and means or IQR for median. Table 2 displays distribution of patient characteristics (row variables) by sex (column variable) which was output after running the code in Fig 1.

It shows factor label and corresponding categories in column 1. Analysis for sex=Male are shown in columns 2-4 and consists of raw or unweighted sample size for each leave of the factor (column 2), weighted column percentages/or median (in column 3) and corresponding 95% CI or IQR (column 4). Analysis for sex=Female are similarly shown in columns 5-7 and displays raw or unweighted sample size for each leave of the factor (column 5), weighted column percentages/or median (in column 6) and corresponding 95% CI or IQR (column 7). Analysis for total Male and Female participants is shown in columns 8-10 and consists of total raw or unweighted sample size for each leave of the factor (column8), weighted total column percentages/or median (in column 9) and corresponding 95% CI or IQR (column 10). To compare the distribution of selected factors by sex, we use the 95% CI or IQR. For instance, among participants aged 40-59, there were more females than males 61.8% (95% CI: 38.2% −85.3%) versus 25.2% (95% CI: 19.9% - 30.5%). This was the reverse among participants aged >= 60 years 62.1% (95% CI: 56.0% - 68.1%) for males versus 19.1% (95% CI: 2.2% - 35.9%) for females. However, median age at screening was comparable at 64.2 years (IQR: 51.3 - 73.0) for males compared to 50.3 years (IQR: 41.1 - 53.8). If missing values were present in the factor variable and the user specifies to display them, then an additional “Missing” category would be to that factor variable.

Table 3 shows the distribution of patient characteristics by hepatitis A test results which was obtained after running the code in Fig 2.

**Fig 2.**
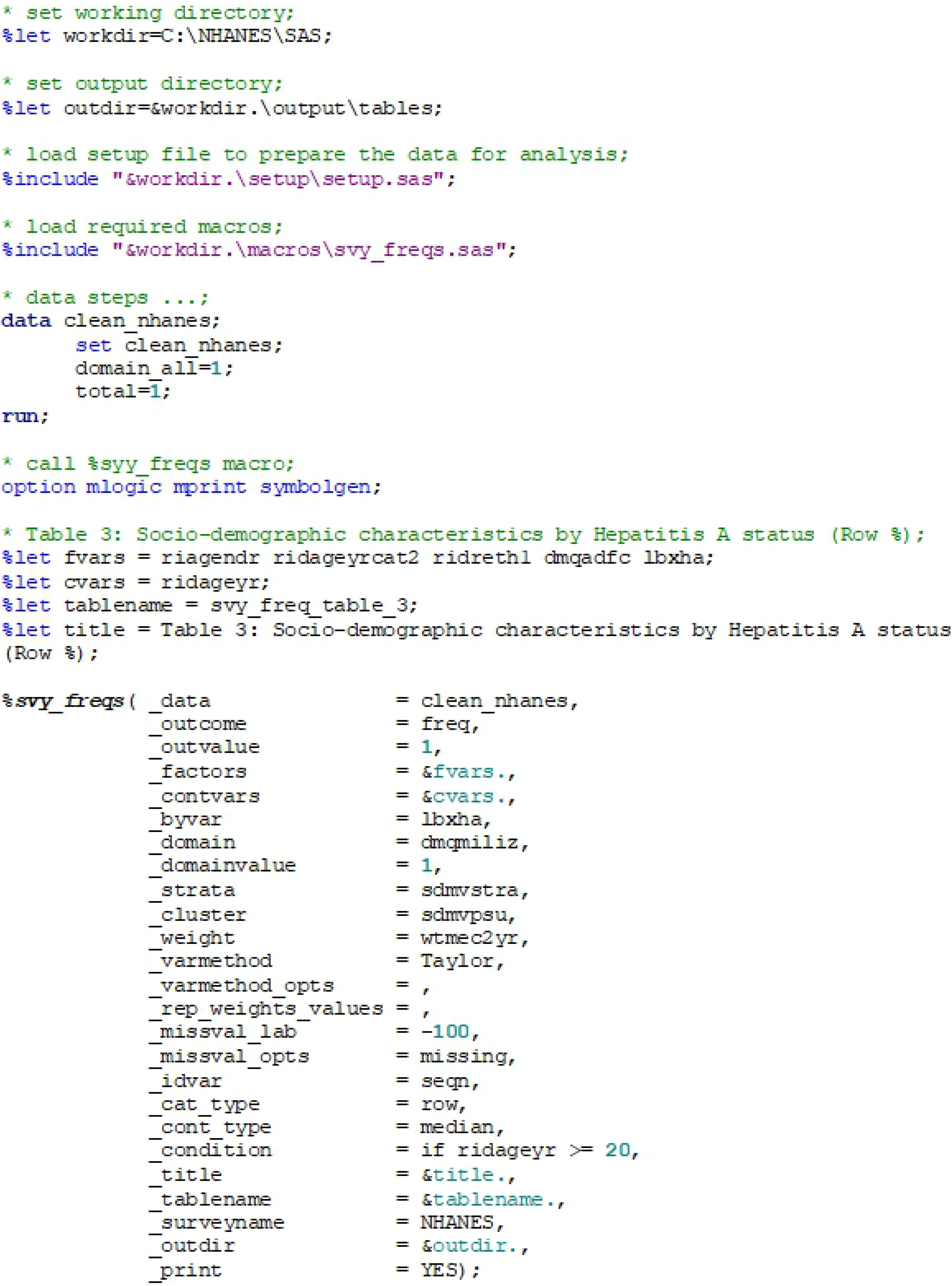
Sample *%svy_freqs* macro call to output row percentages.

The columns are organized in a similar way as described above for Table 2. The output however, presents row percentages and includes columns 8-10 for missing values. The results shows the distribution of hepatitis A status across the given factor variables. It can be seen that there is no significant differences in the distribution of hepatitis A status with 37.1% (95% CI: 33.6 – 40.6) males and 38.9% (95% CI: 17.3 – 60.4) for females reporting positive status. The wider confidence interval for females is attributed to the small sample sizes, n/N=15/36. However, participants aged 20-39 years reported a significantly higher positive hepatitis A status of 82.4% (95% CI: 72.6 - 92.2) compared to 30.8% (95% CI: 23.1 – 38.5) and 30.1% (95% CI: 25.3 – 34.9) for those aged 40-59 and >= 60 years respectively. It is important to note that if missing values are suppressed the estimates will also change since the denominator will have changed.

Table 4 shows distribution of hepatitis A prevalence by sex obtained after running the code in Fig 3.

**Fig 3.**
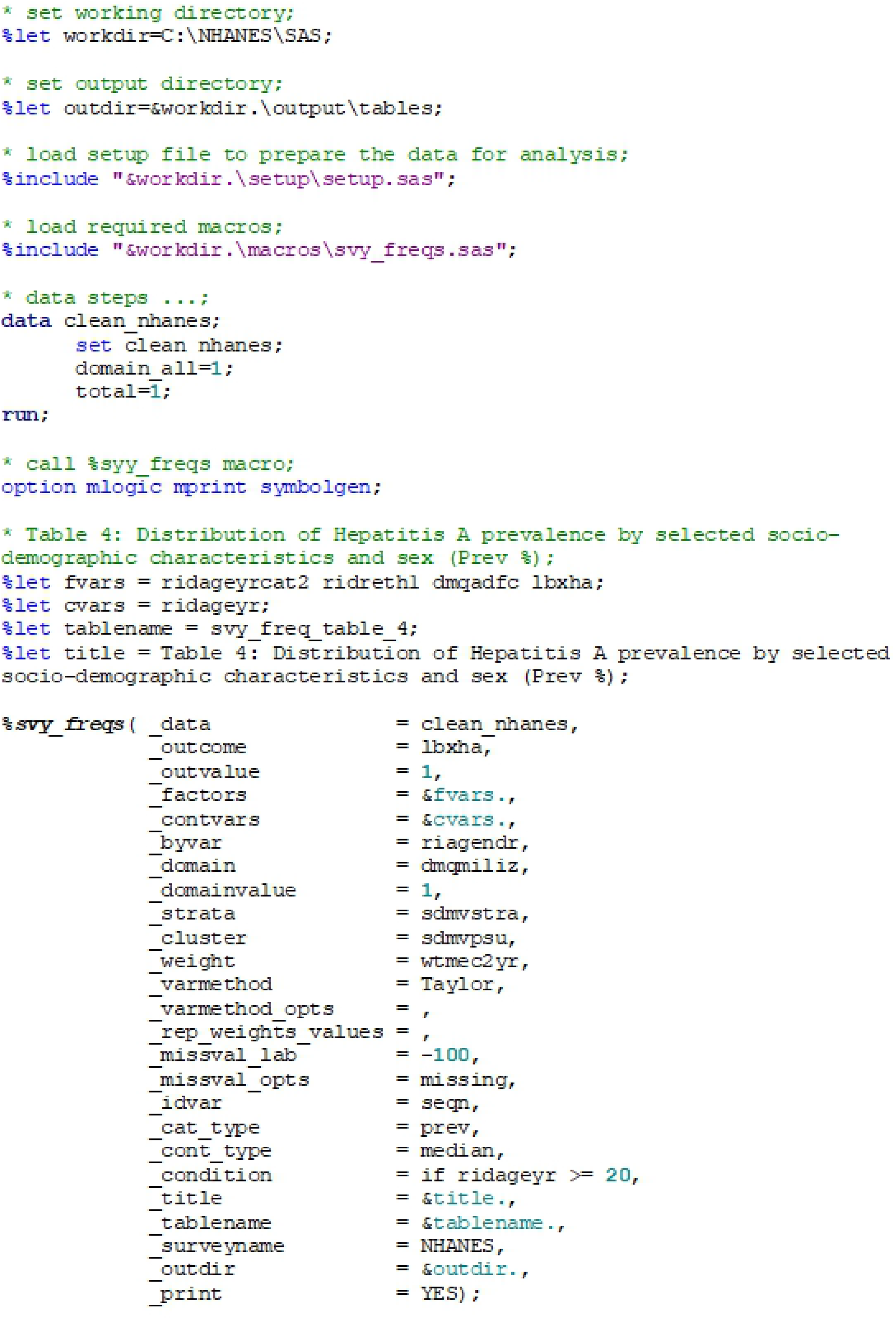
Sample *%svy_freqs* macro call to output prevalence percentages.

The columns are also organized in a similar way as described in previous tables 2 and 3. The output shows there were no significant difference by across each factor by sex as the confidence intervals are overlapping due to the small sample number of females. This serves as a pointer to the reader to use variables that are more balanced as a by-group variable.

## Discussion

This paper presents a simple yet flexible and generalizable SAS macro, ***%svy_freq***, for generating publication-quality outputs from cross-tabulations between a factor and a by-group variable given a third variable using survey or non-survey data. It is also easy to follow, straightforward for the end user and simple for a SAS programmer to extend it. Though it is designed to perform three-way cross-tabulation, as needed for showing the distribution of disease prevalence among one or more factor variables, stratified over a by-group variable, it also performs two-way cross-tabulations similar to those provided by other available macros, resulting in a more comprehensive solution to generating formatted cross-tabulations. It also provides weighted percentages and corresponding 95% confidence intervals. It importantly supports not only the classic domain analyses but also other variance estimation methods including replication-based JK and BRR which are now commonly used with complex survey data.

The macro has several limitations. Firstly, it has been developed under Microsoft windows platform and users might require making appropriate code adjustment to adopt it to other platforms such as Linux. Secondly, it cannot handle arbitrary nesting of by-group variables, such as those supported by PROC TABULATE. Thirdly, it assumes that the user provides a as input data a clean SAS dataset where variables have been labelled and formats for categorical variables defined. Fourthly, it does not provide interpretation of results, so users should consult qualified Statistician for any inference. We however, feel it provides a good tradeoff between simplicity and ease of use, flexibility and generalizability.

## Conclusion

We have also developed another SAS macro, ***%svy_logistic_regression*** [17], for producing publication-quality tables from unadjusted and adjusted logistic regression analyses. We also plan to extend this macro to include other statistical techniques. Our key motivation is to generate tools to automate the process of data analysis which will shorten the time required to prepare output and hence provide quick well-formatted results for consumption and/or dissemination in lieu of reproducible science.

## Supporting information

**S1 Data. Sample NHANES dataset used to demonstrate the functioning of the manuscript.** The complete NHANES dataset is freely available to the public on the NHANES website at: https://www.cdc.gov/nchs/nhanes/Index.htm. (ZIP)

**S2 Code. The SAS macro *% svy_freqs* source code.** The source code for the SAS macro to perform cross-tabulation between a factor and a by-group variable given a third variable and a simple example of the implementation. (TXT).

**S3 Code. Sample code for *% svy_freqs* macro call.** A sample SAS code to read in the data and call the macro. (TXT)

The **S2 Code** and **S3 Code** are also available online at https://github.com/kmuthusi/generic-sas-macros and have been labelled as “%*svy_freqs.sas*” and “*svy freqs anafile.sas*” respectively.

## Funding

This work was supported by the President’s Emergency Plan for AIDS Relief (PEPFAR) through the U.S. Centers for Disease Control and Prevention (CDC). The funders had no role in study design, data collection and analysis, decision to publish, or preparation of the manuscript.

## Acknowledgement

We thank our colleagues in the Surveillance and Epidemiology branch for testing the SAS macro and providing valuable feedback for improvement and for thoroughly reviewing the manuscript.

## Author contribution

JM and SM took part in concept development. JM developed and documented the SAS macro and prepared the final manuscript. SM tested and debugged the SAS macro. PY helped define user requirements and tested the SAS macro. All authors read and approved of the final manuscript for publication.

## Notes

**Conflict of interest:** none

## References

1. StataCorp. Stata: Release 14. Statistical Software. College Station, TX: StataCorp LP; 2015.

2. Watson Ian. tabout: Stata module to export publication quality cross-tabulations: Boston College: MA; 2011. Available from: http://ideas.repec.org/c/boc/bocode/s447101.html.

3. Nguyen Minh TABMULT: Stata module to produce multiple two-way tabulations Statistical Software Components S457371: Boston College: MA; 2012. Available from: https://ideas.repec.org/c/boc/bocode/s457371.html.

4. Sunesara I, Lirette ST, Griswold ME. Survey Tables Binary: A SAS Macro for Publication Quality Tables of Complex Survey Data. Austin Biometrics and Biostatistics. 2015;2(4).

5. Xiong Z. %YAMGAST: Yet Another Macro to Generate a Summary Table. PharmaSUG2008.

6. Martin K. A Researcher’s Guide to Making Descriptive and Analytic Tables that are Ready to Publish. WUSS2006.

7. Zhou Y, Zhang L, Hancock ML. %SummaryTable: A SAS Macro to Produce a Summary Table in Clinical Trial. PharmaSUG2006.

8. Zuo J, Haske CR, editors. Creating Clinical Trial Summary Table Containing P-Values: A Practical Approach Using Standard SAS Macros. SUGI 22.

9. Bhaskar B, Murray K. Generating Customized Analytical Reports from SAS Procedure Output.

10. Arnold T, Kuhfeld WF. Using SAS and LATEX to Create Documents with Reproducible Results. URL: http://supportsascom/resources/papers/proceedings12/324-2012pdf. 2012.

11. Peng RD, Dominici F, Zeger SL. Reproducible epidemiologic research. American Journal of Epidemiology. 2006:163(9):783–9. doi: 10.1093/aje/kwj093. PubMed PMID: 16510544.

12. Peng RD. Reproducible research and Biostatistics. Biostatistics. 2009:10(3):405–8. doi: 10.1093/biostatistics/kxp014. PubMed PMID: 19535325.

13. Peng RD. Reproducible research in computational science. Science. 2011:334(6060):1226–7. doi: 10.1126/science.1213847. PubMed PMID: 22144613: PubMed Central PMCID: PMCPMC3383002.

14. Patil P, Peng RD, Leek JT. What Should Researchers Expect When They Replicate Studies? A Statistical View of Replicability in Psychological Science. Perspectives on Psychological Science. 2016:11(4):539–44. doi: 10.1177/1745691616646366. PubMed PMID: 27474140; PubMed Central PMCID: PMCPMC4968573.

15. Levy PS, Lemeshow S. Sampling of Populations: Methods and Applications. 4th ed. New York: John Wiley & Sons; 2013.

16. Lohr SL. Sampling: Design and Analysis. 2nd edition ed. Boston: Brooks/Cole; 2010.

17. Muthusi J, Mwalili S, Young P. %svy_logistic_regression: A generic SAS macro for simple and multiple logistic regression and creating quality publication-ready tables using survey or non-survey data. PLoS One. 2019:14(9):e0214262. Epub 2019/09/04. doi: 10.1371/journal.pone.0214262. PubMed PMID: 31479445.

18. Cochran W. Sampling Techniques. 3rd ed. New York: John Wiley & Sons; 1977.

19. Kish L. Survey Sampling. New York: John Wiley & Sons; 1965.

20. Foreman E. Survey Sampling Principles. New York: Marcel Dekker; 1991.

21. SAS Institute Inc. SAS/STAT® 9.3 User’s Guide. Cary, NC: SAS Institute Inc; 2011.

22. SAS Institute Inc. Base SAS® 9.3. Cary, NC: SAS Institute Inc; 2011.

23. Wolter KM. Introduction to Variance Estimation. New York: Springer-Verlag; 1985.

24. Kalton G, Kasprzyk D. The Treatment of Missing Survey Data. Survey Methodology. 1986:12:1–16.

25. Cochran WG. Sampling Techniques. 3rd Edition ed. New York: John Wiley & Sons.; 1977.

26. Brick JM, Kalton G. Handling Missing Data in Survey Research. Statistical Methods in Medical Research. 1996:5:215–38.

27. NASCOP. National AIDS and STI Control Programme (NASCOP), Kenya. Kenya AIDS Indicator Survey 2012: Final Report. Nairobi. 2014.

28. NASCOP. National AIDS/STI Control Programme (NASCOP), Kenya. 2007 Kenya AIDS Indicator Survey: Final Report. Nairobi. 2009.

29. KNBS, MOH, NASCOP, KEMRI, NCPD, ICF International. Kenya National Bureau of Statistics, Ministry of Health/Kenya, National AIDS Control Council/Kenya, Kenya Medical Research Institute, National Council for Population and Development/Kenya, and ICF International. Kenya Demographic and Health Survey 2014. Rockville, MD, USA. 2015.

30. KNBS, ICF Macro. Kenya National Bureau of Statistics (KNBS) and ICF Macro. Kenya Demographic and Health Survey 2008-09. Calverton, Maryland. 2010.

31. CBS, MOH, ORC Macro. Central Bureau of Statistics (CBS) [Kenya], Ministry of Health (MOH) [Kenya], and ORC Macro. Kenya Demographic and Health Survey 2003. Calverton, Maryland. 2004.

32. Johnson CL, Dohrmann SM, Burt VL, Mohadjer LK. National Health and Nutrition Examination Survey: Sample design, 2011–2014. National Center for Health Statistics. Vital and Health Statistics. 2014:2(162).

33. Centers for Disease Control and Prevention. Centers for Disease Control and Prevention (CDC). National Center for Health Statistics (NCHS). National Health and Nutrition Examination Survey Data. Hyattsville, MD: U.S. Department of Health and Human Services, Centers for Disease Control and Prevention, 2013-2014, URL: https://www.cdc.gov/nchs/nhanes/Index.htm: National Center for Health Statistics (NCHS); 2015.

